# Whole Chloroplast Genomes reveals the uniqueness of Bolivian native cacao (*Theobroma cacao*) from the northern part of Bolivia

**DOI:** 10.1101/2021.04.16.440153

**Authors:** M Gumiel, OM Rollano-Peñaloza, C Peralta-Rivero, L Tejeda, V Palma, P Cartagena, P Mollinedo, JM Peñarrieta

## Abstract

We report the complete chloroplast sequences of two varieties of *Theobroma cacao* collected in the Bolivian Amazonia using Next-Generation Sequencing. Comparisons made between these two chloroplast genomes and the Belizean reference plastid genome identified 19 and 22 nucleotide variants. The phylogenetic analysis reported three main *T. cacao* clades belonging to the Forastero, Criollo and Trinitario groups. The Bolivian Native Cacao varieties were located inside the Trinitario group forming their unique branch. The Bolivian Native Cacao branch reveals a possible new subpopulation different from the well-characterized *T. cacao* subpopulations. The phylogenetic trees showed that the relationships among the *T. cacao* varieties were consistent with their geographical locations placing the Cacao Center of Origin in Western Amazon. The data presented here will contribute to the usage of ultrabarcoding to distinguish different *T. cacao* varieties and to identify native cacaos from introduced cacaos. Thus helping in the conservation of local native varieties of *T. cacao*.

## INTRODUCTION

*Theobroma cacao* is a tree cultivated in tropical and subtropical regions of the world. In Bolivia, *T. cacao* is cultivated in the humid regions of Beni, Pando, La Paz, Cochabamba and Santa Cruz departments and is a source of economical sustainability for families. The final product of the processed cacao is chocolate, cocoa butter or cocoa powder. In Bolivia as well as other Latin American countries, numerous commercial cacao varieties were introduced by the governments, and mixed with the Bolivian native cacao varieties endangering such native species (Bazoberry Chali and Salazar Carrasco, 2008). The high biodiversity of cacao varieties found in Bolivia is little studied, and therefore more research efforts needs to be done. The main interest lies into characterize and identify the phylogenetic relationships between these subpopulations and aid in the accurate subspecies identification.

The classification of *T. cacao* has been traditionally divided in: Criollo, Forastero and Trinitario (Cheesman, 1944). The product derived from the varieties corresponding to the Criollo is considered of being the best quality, whereas the Forastero varieties present phenotypes that are more disease resistant. On the other hand, Trinitario varieties are supposed to be originated from hybridization between Criollo and Forastero groups, and presents the best characteristics of both lineages. Vast genetic analyses using different molecular markers revealed a huge number of genetic groups (de la Cruz et al., 1995) localized in the Amazonia region and in Central America. Motamayor et al. (2008) proposed ten genetically differentiated groups: Amelonado, Contamana, Criollo, Curaray, Guianna, Iquitos, Marañon, Nacional, Nanay and Purus. Cornejo et al. (2018), corroborated with a population genomic analyses that these ten groups underwent strong domestication. Moreover, they found that the Criollo, Amelonado and Nacional varieties contributed to the individual ancestry, and that the samples of the Amazonian Basin present a higher diversity in contrast to the lower diversity found in the samples from the Atlantic side.

The Bolivian native cacao varieties have not being considered within the ten groups described above (Motamayor et al., 2008; Cornejo et al., 2018) and they might be a different population. These studies have been done with Population Genetics but could also be resolved with Barcoding. Nowadays, as the DNA sequencing prices have gone notably down is more accessible to sequence entire genomes or whole plastid genome rather than genes or DNA fragments. Whole plastid genomes as the chloroplast genome are more conserved than the nuclear genome. Plastid genomes enclose essential information markers for phylogenetic relationships among closely related taxa due to the low rate of polymorphism in the chloroplast. Therefore whole chloroplast genome sequencing has become an interesting barcoding tool for plants called ultra-barcoding (Kane and Cronk, 2008). Ultra-barcoding is based on data derived from high-throughput sequencing also called Next-Generation Sequencing (NGS). NGS data is more sensitive than traditional molecular markers (e.g. microsatellite) because genome target regions are significatively larger.

In this work, we focus on *T. cacao* ultra-barcoding which was used to identify genetic variation below species level. The results obtained in this work will provide valuable DNA sequence information for taxonomic studies and the development of molecular markers for below-species level identification of *T. cacao* coming from Bolivia. Ultimately, providing tools for *T. cacao* germplasm conservation.

## MATERIALS AND METHODS

### Plant material

*Theobroma cacao* fully developed leaves were collected, from two different varieties in two different regions from Bolivia: “mir20” from an indigenous Community (Miraflores, Pando) and “naz7” from a peasant community (Nazareth, Beni). Each tree received standard agronomic practices, and the cacao samples gained prizes for the best chocolate product in 2013, 2015, 2017 and 2019 at the international contest “Salon du Chocolat” in Paris, France. The leaves were collected and dry-stored at −20°C.

### DNA isolation and sequencing

5 mg of frozen leaves were crushed in a mortar to obtain fine powder using liquid nitrogen and the powder was transferred to microcentrifuge tubes. DNA isolation was performed using the Purelink Genomic DNA Kit (Thermo, CA, USA) according to the procedure described in the manufacture’
ss instructions. Chloroplast genome sequencing was outsourced to Omega Bioservices, USA and sequenced on a Miseq (Illumina, CA, USA). Using 2 x 150 bp paired-end reads generated with the Nextera Truseq libraries (Illumina, CA, USA).

### Bioinformatic analysis

Reads were trimmed and cleaned Assembly, mapping and short read post-processing were performed using Velvet (1.2.10), Bowtie2 (bowtie-bio.sourceforge.net/bowtie2, V. 2.3.5.) and Samtools utilities (htslib.org/). The annotation of the chloroplast genes was made by GeSeq (Tillich et al., 2017). A chlorogenome map was generated using OGDraw (Greiner, Lehwark, and Bock, 2019).

## RESULTS

Trimmed and cleaned reads were further filtered out by mapping them against a *Theobroma cacao* reference chloroplast genome (RefSeq assembly: GCF_000208745.1, National Center for Biotechnology Information). The chloroplast reference genome corresponds to the sample Scavina-6 from Perú (Kane et al., 2012; Argout et al., 2017). A total of approximately 47 million reads (mir20 sample) and 25 million reads (naz7 sample) were used in the assembly of the two plastid genomes (Table 1). The chloroplast reference genome (HQ244500.2) has 160,619 base pairs and our plastid coverage was more than a 100X for both samples. The coverage was enough to assemble the plastid genomes of both samples. The two plastids from the *T. cacao* Bolivian varieties differ in only 19 nucleotides for mir20 and 22 nucleotides for naz7 compared to the Belizean plastid genome. The two plastid genomes were deposited in GenBank under accessions: MW243993 for mir20 and MW243994 for naz7) The GC content of both samples was 36,9 % (Figure 1).

**Table 1.**
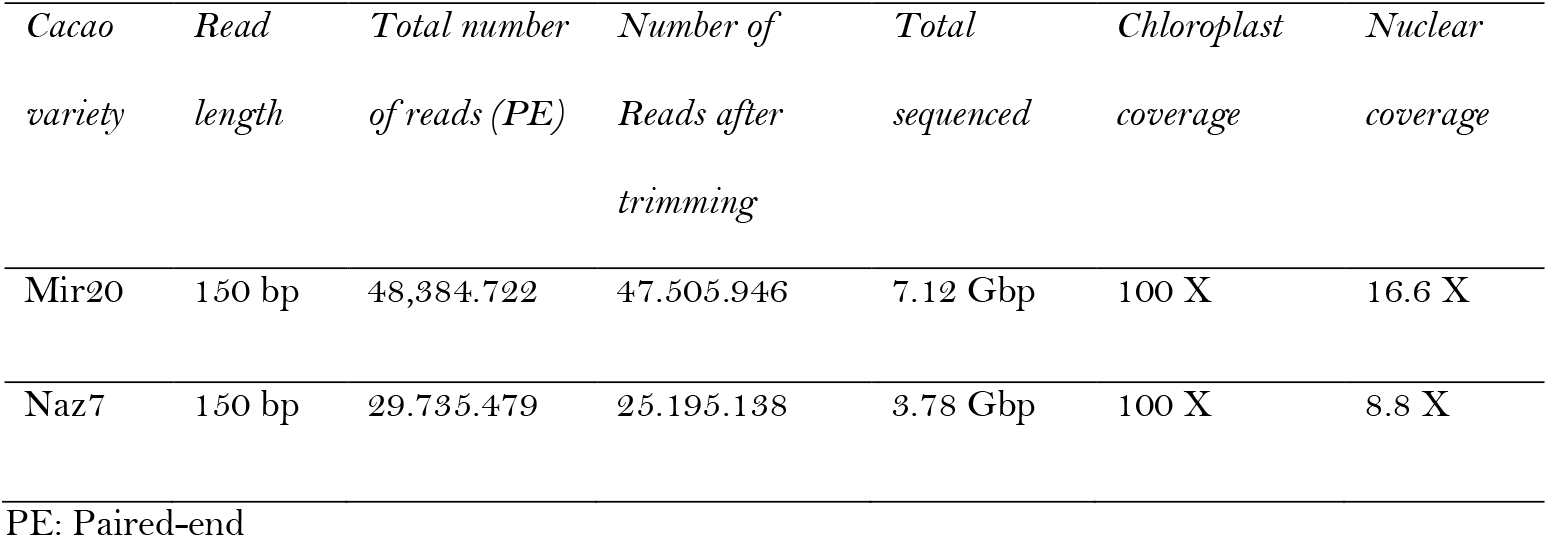
Illumina sequence summary statistics and observed coverage of the nuclear and chloroplast genome for Bolivian native cacao varieties.

**Fig 1.**
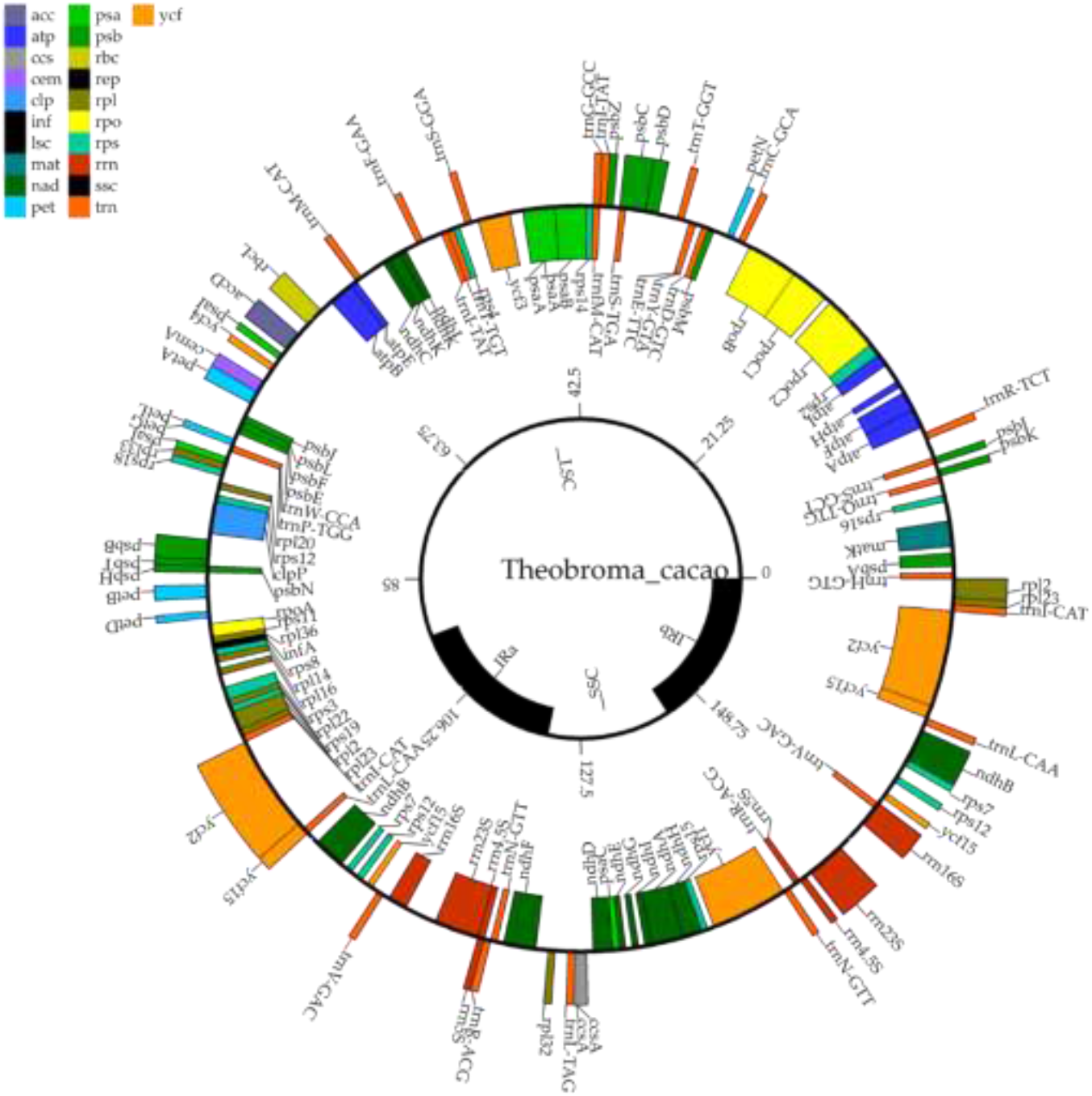
*Map of Theobroma cacao var. naz7 chloroplast genome. The thick lines indicate the extent of the two inverted repeat regions (IRa and IRb), which separate the genome into the small single-copy (SSC) and the large single-copy (LSC) region. Colors indicate different gene functional groups*. Abbreviations: *acc*: Acetyl-CoA carboxylase, *atp*: Subunits of ATP synthase, *ccs*: c-type cytochrome synthesis gene, *cem*: Envelope membrane protein, *clp*: Protease, *inf*: Translation initiation factor, *mat*: Maturase, *nad*: Subunits of NADH dehydrogenase, *pet*: Subunits of cytochrome b/f complex, *psa*: Subunits of photosystem I, *psb*: Subunits of photosystem II, rbc: Large subunit of rubisco, *rep*: repeated regions, *rpl*: Proteins of large ribosomal subunit, *rpo*: Subunits of RNA polymerase, *rps*: Proteins of small ribosomal subunit, *rrn*: Ribosomal RNAs, *trn*: Transfer RNAs, *ycf*: Conserved hypothetical chloroplast reading frames.

The comparison between the Belizean reference plastid genome and the Mir20 sample identified 19 different nucleotides. Comparing the reference plastid genome with the Naz7 sample 22 variant nucleotide positions were observed. The Bolivian cacao plastid annotation contained 132 genes including 8 ribosomal RNA, 37 tRNA genes and 85 protein-coding genes.

To explore the phylogenetic relationships we included 13 plastid genomes in our analysis (Table 2). The phylogenetic analyses revealed significant divergence between clades of *T. cacao* from the diverse varieties. The Maximum Likelihood tree showed two strongly supported *T. cacao* clades. The clades on both ends correspond to the Forastero and Criollo Groups. The clade in the middle belongs to the hybrid group between them, the Trinitario group (Fig. 2). The Bolivian Native Cacao forms a different group inside the hybrid clade. The tree also verifies that the Trinitario varieties (e.g. ICS01, ICS06) are hybrids between Forastero and Criollo. The *T. cacao* plastid genome accession HQ336404.2 (Jansen et al., 2011), which has no publicly available information about its geographical origin groups strongly with the Forastero plastid variety (e.g. Amelonado) (Fig. 2).

**Table 2.** List of the 13 chloroplast genomes used in this study. The accessions from the National Center for Biotechnology Information (NCBI) are listed for easy reference..

| Variety | Genbank Accession | Origin | Groups according to our<br>barcoding analysis | Traditional variety<br>classification |
| --- | --- | --- | --- | --- |
| Amelonado | JQ228380.1 | USDA, TRAS, Puerto Rico . | Forastero | Forastero<sup>1</sup> |
| - | HQ336404.2 | - | Forastero | Unknown<sup>2</sup> |
| - | JQ228389.1 | GI328924764 | Forastero | Unknown<sup>1</sup> |
| Scavina-6 | HQ244500.2* | Peru | Unknown | Forastero<sup>1</sup> |
| ICS-01 | JQ228381.1 | USDA, TARS, Puerto Rico | Trinitario? | Trinitario<sup>1</sup> |
| ICS-06 | JQ228383.1 | USDA, TARS, Puerto Rico | Trinitario? | Trinitario<sup>1</sup> |
| Scavina 6 | NC_014676.2 | Peru | Unknown | Forastero<sup>1</sup> |
| Scavina-6 | JQ228382.1 | Peru | Unknown | Forastero<sup>1</sup> |
| Stahel | JQ228385.1 | Suriname | Criollo | Trinitario<sup>1</sup> |
| Criollo 22 | JQ228379.1 | USDA, SPCL, Beltsville, MD | Criollo | Criollo<sup>1</sup> |
| EET-64 | JQ228384.1 | USDA, TARS, Puerto Rico | Criollo | Forastero – Trinitario Hybrid<sup>1</sup> |
| ICS-39 | JQ228387.1 | USDA, TARS, Puerto Rico | Criollo | Trinitario<sup>1</sup> |
| Pentagonum | JQ228386.1 | USDA, TARS, Puerto Rico | Criollo | Criollo<sup>1</sup> |
| Mir20 | MW243993 | Bolivia | Bolivian Native Cacao | Unknown<sup>3</sup> |
| Naz7 | MW243994 | Bolivia | Bolivian Native Cacao | Unknown<sup>3</sup> |

\*Reference plastid Genome
1. Kane et al., 2012
2. Jansen et al., 2010
3. This paper

**Fig 2.**
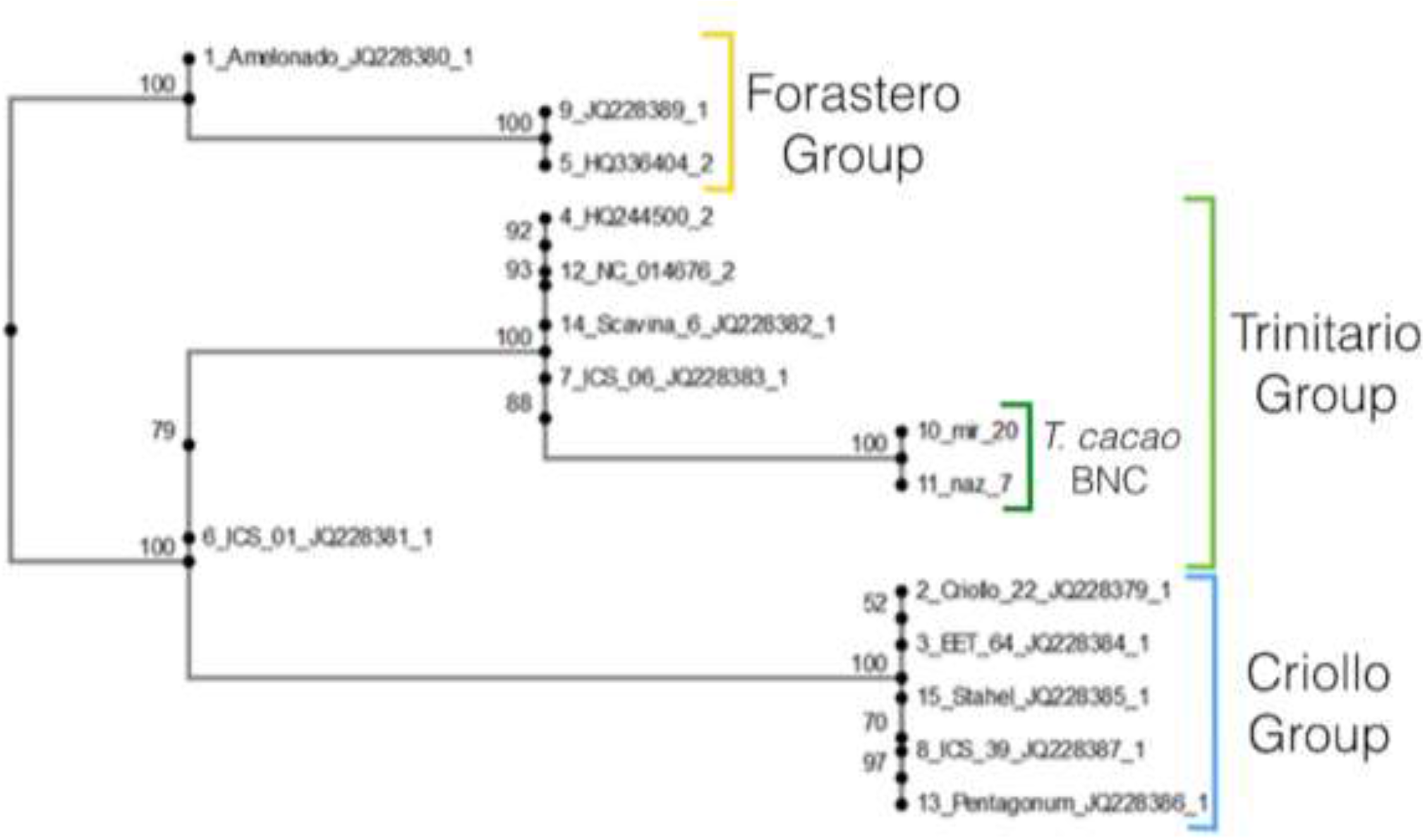
The cacao phylogenetic tree constructed with chloroplast genomes reveals a unique group formed by Bolivian native cacao (BNC) varieties. Bootstrap values are given in percentage (%). Details of the chloroplast genome accessions are in Table 2.

## DISCUSSION

DNA barcoding is useful for several applications including identification below species level, estimating phylogenetic diversity and to identify species that are new to science. DNA barcoding allows to identify taxon through DNA sequencing using specific nuclear locus (e.g. ITS region) (Bellemain et al., 2010), mitochondrial genes (e.g. *COI and CytB*) (Degli Esposti et al., 1993; Hebert et al., 2003) or complete genomes. The results reported in this work shows the benefits of whole chloroplast genome barcoding for organism identification below the species level as the Bolivian Native Cacao is identified as a different group from other *T. cacao* varieties (Fig. 2).

The *T. cacao* phylogenetic tree constructed with whole chloroplast genomes revealed three different clades below the species level (Fig. 2). The clades formed for the Criollo and Forastero group have been reported before with whole chloroplast genome sequencing as a barcoding tool (Kane et al., 2012). The Forastero group represents the cluster of cacaos from South America and the Criollo group represents the Central America group. A third clade was observed in our study and was formed mostly by the Trinitario hybrid variety (Fig. 2). This clade includes the Bolivian samples in a separate branch. The Bolivian Native Cacao forming a different cacao subpopulation has been already reported with microsatellites (Zhang et al., 2012). Bolivian Native Cacao is very likely that will form a new subgroup among the ten subpopulations described by Cornejo et al. (2018) and Motamayor et al. (2008).

The fact that Bolivian Native Cacao samples are inside the Trinitario group might be explained because most of the *T. cacao* varieties in this group live in the Western Amazon and the pacific coast of South America, just in the middle of the Criollo distribution zone (Central America) and the Forastero distribution zone (Atlantic coast of South America). The Western Amazon has been proposed as the Center of Origin for Cacao species through population genomics (Cornejo et al. 2018) and archeological evidence (Zarrillo et al., 2018). Thus, we suggest that the Trinitario group should be renamed as the Center of Cacao Origin group. Thus, avoiding the misconception that many of these varieties are hybrids between Forastero and Criollo.

## CONCLUSIONS

Whole chloroplast genome sequencing of *T. cacao* is a useful approach to identify cacao varieties below the species level. The sequencing of more cacao varieties will deepen the results obtained in this work, showing the Center of Cacao Origin in the Western Amazon also through Ultrabarcoding.

## ACKNOWLEDGEMENTS

The authors would like to thankfully acknowledge the support of Centro de Investigación y Promoción del Campesinado (CIPCA), the Bolivian IDH funding at Universidad Mayor de San Andrés and the Swedish International Development Cooperation Agency (SIDA). The authors would like to express their gratitude to cacao-producing communities in Beni and Pando, Bolivia for his enormous help and support in this study.

## AUTHOR CONTRIBUTIONS

PC, JMP, PM designed the study. MG, CP, OMRP, conducted the analysis, data interpretation and drafted the manuscript. LT and VP conducted part of the analysis and experiments. PC, JMP, PM, and CP supervised the work. All the authors contributed to and approved the final manuscript.

